# Starless bias and parameter-estimation bias in the likelihood-based phylogenetic method

**DOI:** 10.1101/435412

**Authors:** Xuhua Xia

## Abstract

I analyzed various site pattern combinations in a 4-OTU case to identify sources of starless bias and parameter-estimation bias in likelihood-based phylogenetic methods, and reported three significant contributions. First, the likelihood method is odd in that it may not generate a star tree with sequences that are equidistant from each other. This behaviour, dubbed starless bias, happens in a 4-OTU tree when there is an excess (i.e., more than expected from a star tree and a substitution model) of conflicting phylogenetic signals supporting the three resolved topologies equally. Special site pattern combinations leading to rejection of a star tree, when sequences are equidistant from each other, were identified. Second, fitting gamma distribution to model rate heterogeneity over sites is strongly confounded with tree topology, especially in conjunction with the starless bias. I present examples to show dramatic differences in the estimated shape parameter α between a star tree and a resolved tree. There may be no rate heterogeneity over sites (with the estimated α > 10000) when a star tree is imposed, but α < 1 (suggesting strong rate heterogeneity over sites) when an (incorrect) resolved tree is imposed. Thus, the dependence of “rate heterogeneity” on tree topology implies that “rate heterogeneity” is not a sequence-specific feature, cautioning against interpreting a small α to mean that some sites are under strong purifying selection and others not. Thirdly, because there is no existing (and working) likelihood method for evaluating a star tree with continuous gamma-distributed rate, I have implemented the method for JC69 in a self-contained R script for a four-OTU tree (star or resolved), in addition to another R script assuming a constant rate over sites. These R scripts should be useful for teaching and exploring likelihood methods in phylogenetics.

## 1. INTRODUCTION

I explore two phylogenetic issues here. The first on the starless bias. If a set of aligned sequences are equidistant from each other, i.e., the number of various types of substitutions between any two sequences is exactly the same, then we intuitively would expect a star tree. Distance-based methods will indeed give us a star tree whenever pairwise distances are all equal. The starless bias refers to the inability of a phylogenetic method to generate a star tree with equidistant sequences. It was first alluded to in a study of potential bias in maximum likelihood method involving missing data and rate heterogeneity over sites (Xia, 2014), but its occurrence is more general that that. I will illustrate this bias here with four sequences, identify the source of the bias, and discuss its relevance to the star-tree paradox associated with Bayesian phylogenetic inference (Lewis et al., 2005; Yang, 2007a; Yang and Zhu, 2018).

The second issue, related to the first, is the confounding effect of tree topology on phylogenetic parameter estimation, in particular the shape parameter α of gamma distribution used to model rate heterogeneity over sites. It may seem obvious that α depends on topology, but the issue needs to be studied for two reasons. First, how such dependence occurs is not well dissected. Second, one frequently encounters interpretation of α as if it is a sequence-specific feature, with a small alpha interpreted as indicating some sites strongly constrained by purifying selection and other sites not. It is therefore relevant to caution against such interpretation with real examples. For simplicity, I will work on 4-OTU trees only so that all site patterns can be conveniently considered.

I provide two self-contained R script files (New2.R and NewGamma3.R) implementing the likelihood method with JC69 model (Jukes and Cantor, 1969) with four OTUs. New2.R assumes a constant rate over sites and estimates branch lengths for the star tree (with four branches) and a resolved tree (with five branches). NewGamma3.R does the same but with a continuous gamma-distributed rate over sites. They are plain text files that can be copied and pasted into an R window to generate results presented in the paper, and should also be useful for teaching and exploring likelihood methods in phylogenetics.

### Site patterns classification and sequence representation

With four sequences, there are 256 possible site patterns. For sequences to evolve according to the JC69 substitution model (Jukes and Cantor, 1969), the 256 site patterns would become equally frequently after sequences experienced full substitution saturation. Different combinations of these 256 site patterns support different substitution models and different topologies. We will focus on the JC69 substitution model and classify these 256 possible site patterns into 15 classes (Fig. 1A) so that we only need to calculate likelihood for these 15 site pattern classes. The transition probabilities needed for likelihood calculation for the JC69 model, together with other frequently used Markovian nucleotide substitution models such as F84 (used in DNAML since 1984, Hasegawa and Kishino, 1989; Kishino and Hasegawa, 1989), HKY85 (Hasegawa et al., 1985), TN93 (Tamura and Nei, 1993), and GTR (Lanave et al., 1984; Tavaré, 1986) have been numerically illustrated and re-derived with three different approaches (Xia, 2017; Xia, 2018c). These illustrations are sufficiently detailed for one to extend the JC69 model in the two R script files to other models.

**Fig. 1.**
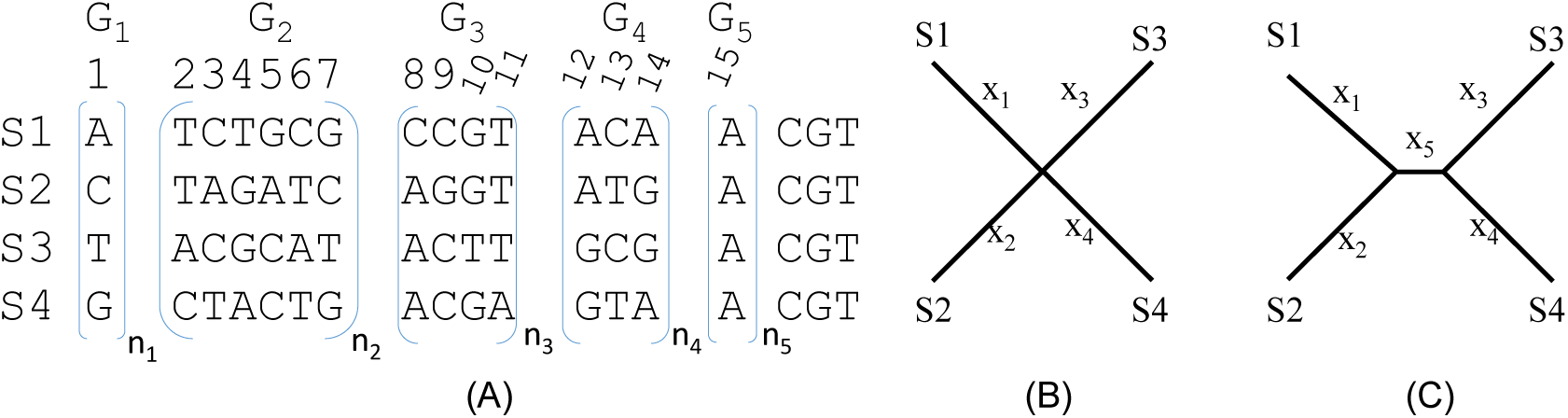
Representative site patterns and alternative trees used. (A) A total of 256 possible site patterns with four sequences, classed into 15 site patterns relevant to the JC69 model and further boxed into five site pattern groups (G_1_ to G_5_). Sites within each group jointly do not support any of the three alternative resolved trees. (B) A star tree. (C) One of three resolved trees with x_5_ > 0.

The 15 site pattern classes (Fig. 1A) can be further lumped into five groups (G_1_ to G_5_, Fig. 1A). Sites in G_1_ have all four OTUs (S1 to S4) with different nucleotides, with 24 unique site patterns represented as a single G_1_ site in Fig. 1A (because JC69 sees these 24 site patterns as identical). Sites in G_2_ each has three different nucleotides, with a total of 144 unique site patterns. Only six sites are used to represent them in Fig. 1A because JC69 sees these 144 unique sites in this group to be identical to one of the six representative sites. Sites in G_3_ feature two nucleotides, with three OTUs having the same nucleotide, and a total of 48 unique sites represented by four sites in Fig. 1A. Sites in G_4_ also feature two nucleotides, with two OTUs sharing one nucleotide and the other two OTUs sharing the other (i.e., they are the traditional informative sites in Fitch parsimony). There are 36 unique G_4_ sites represented by three sites in Fig. 1A. G_5_ sites are monomorphic, with four unique site patterns represented by site 15 in Fig. 1A.

Given a JC69 model, the five groups of sites (G_1_ to G_5_ in Fig. 1A) support the three unrooted topologies equally (i.e., they do not preferentially support any of the three). This is obvious for G_1_ and G_5_ sites. The six sites in G_2_, shown in Fig. 1A, jointly also support the three topologies equally, so do the four sites in G_3_ and three sites in G_4_. Note that, with a star tree and JC69, we cannot have G_1_, G_2_ and G_4_ sites without having G_3_ sites first. With low sequence divergence, almost all site patterns from a star tree should be G_3_ sites.

Subscripts n_1_ to n_5_ in Fig. 1A mean multiples of the enclosed sites, e.g., an n_2_ of 10 means that the six G_2_ sites in Fig. 1A is repeated 10 times (for a total of 60 sites). A combination of (n_1_, n_2_, n_3_, n_4_, n_5_), where n_i_ corresponds to those in Fig. 1A, means a set of aligned sequences containing n_1_ G_1_ sites, n_2_ G_2_ sites (i.e., n_2_*6 sites), n_3_ G_3_ sites (for a total of n_3_*4 sites), n_4_ G4 sites (for a total of n_4_*3 sites), and n_5_ G5 sites (Fig. 1A). A set of four sequences of length 256 containing all 256 possible site patterns is specified as (24, 24, 12, 12, 4). Such a set of sequences will naturally have equal nucleotide frequencies and twice as many transversions as transitions, i.e., the equilibrium ratio of substitution saturation. A set of sequences with any combinations of (n_1_, n_2_, n_3_, n_4_, n_5_) are equidistant from each other from a JC69 perspective, and we would desire to have a star tree (Fig. 1B) instead of one of the three resolved trees (Fig. 1C). Hereafter I specify a set of four aligned sequences equidistant form each other simply by (n_1_, n_2_, n_3_, n_4_, n_5_) which guarantee that the four sequences are equidistant from each other.

Not all (n_1_, n_2_, n_3_, n_4_, n_5_) combinations are equally likely under the JC69 model with a star tree. For example, the site pattern combination (24,24,12,96,32) has a near-zero chance to occur with a star tree and a strict JC69 model, because, given a star tree with x_5_ = 0, G_4_ sites can only emerge through independent substitutions along each of the four branches (i.e., all G_4_ sites result from convergent substitutions). This means that we cannot have G_4_ sites in a star tree without first having G_3_ sites, so G_4_ sites should not be more frequent than G_3_ sites given a star tree and JC69. The ratio of G_3_/G_4_ sites will decrease from ∞ (when the first substitution occurs) towards 4/3 (i.e., 48 possible G_4_ site patterns and 36 possible site patterns) when sequences have gradually experienced full substitution saturation. The site pattern combination above with a ratio of G_3_/G_4_ equal to (12*4)/(96*3) cannot happen with a star tree because the ratio is far too small.

However, when different genes evolving under JC69 models with different rates (and potentially with different evolutionary histories and conflicting phylogenetic signals) are concatenated, strange site pattern combinations such as (24, 24, 12, 96, 32) may occur and cannot be dismissed as unreal. In fact, this site pattern combination results from a concatenation of site patterns from simulations with four different trees, i.e., the star tree and the three resolved trees (all with JC69 with no rate heterogeneity over sites), with slight modifications to ensure 1) that the nucleotide frequencies are exactly equal to 0.25, 2) that the number of transversions is exactly trice as many as the number of transitions, and 3) that the number of transitional and transversional differences between each pairwise comparison is exactly the same.

### The starless bias

The site pattern combination (24, 24, 12, 96, 32) mentioned above represents a special deviation from any of the four topologies (the star tree plus the three resolved trees). While the four sequences will remain equidistant from each other with the site pattern combination of (24,24,12,96,32), the likelihood method will not favour a star tree in spite of equidistance among sequences. In general, as illustrated in Fig. 2, we expect that increasing number of G_4_ sites relative to G_3_ sites would increase the likelihood of rejecting a star tree. As I mentioned before, G_3_ sites are the first site pattern to appear in sequence evolution along a star tree. In contrast, a G_4_ site requires a minimum of two substitutions when x_5_ = 0 (Fig. 2C), but only a minimum of one with a resolved tree from a parsimony perspective (Fig. 2D). Tree lnL would be greater with x_5_ > 0 (so that a single substitution can occur along the internal branch to be shared by two descendants) than with x_5_ = 0 (which would force two independent substitutions). Thus, G_4_ sites favor a resolved tree (x_5_ > 0). In short, increasing number of G_4_ sites relative to G_3_ sites increases the likelihood of rejecting the star tree. This is true even with G_4_ sites support all three resolved topologies equally because 1/3 of the G_4_ sites will support a resolved topology.

**Fig. 2.**
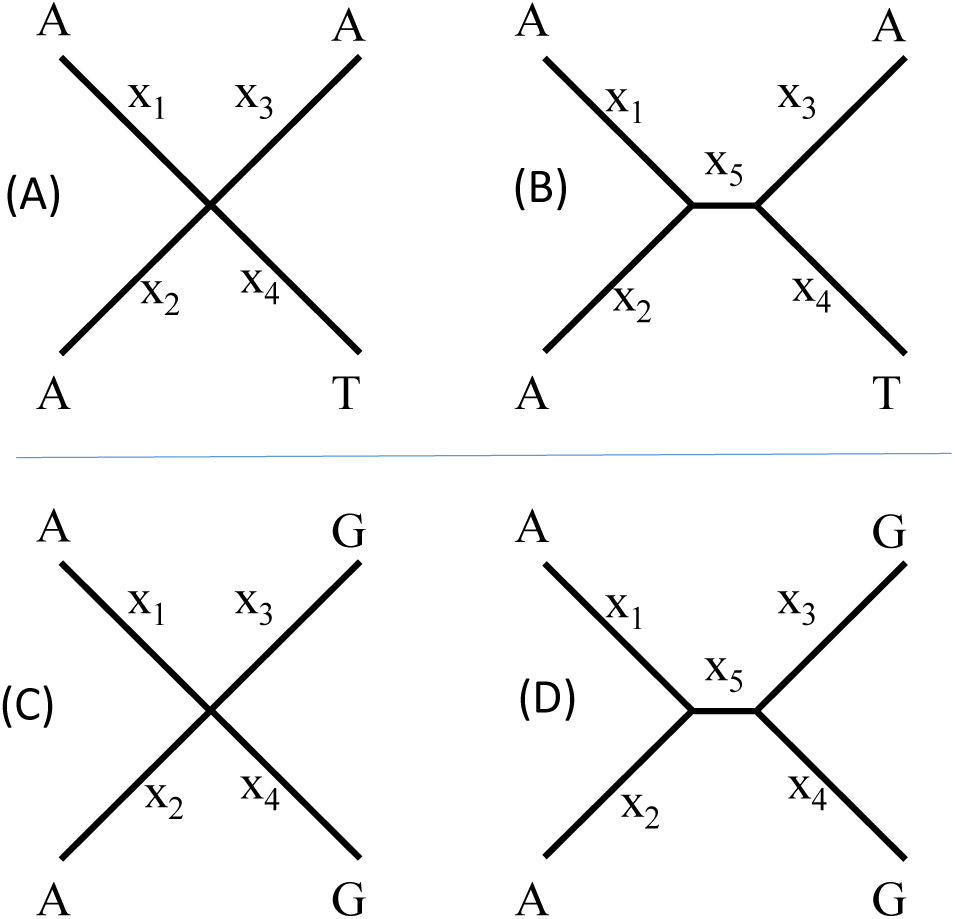
G_3_ and G_4_ sites support star tree and resolved tree differently. (A) and (B) a G_3_ site mapped to a star tree and a resolved tree, respectively. (C) and (D) a G_4_ site mapped to a star tree and a resolved tree, respectively.

### Characterization of rate heterogeneity is confounded by topology

I will use G_4_ sites to illustrate the effect of topology on rate heterogeneity over sites. With a star tree in Fig. 2C, all G_4_ sites require exactly the same number of changes (a minimum of two substitutions per site from a parsimony perspective). In other words, there is no rate heterogeneity among G_4_ sites with a star tree. However, for a resolved tree in Fig. 2D, 1/3 of the G_4_ sites (with identical nucleotides between sister groups as in Fig. 2D) would require only one substitution from the parsimony perspective. The other 2/3 of the G_4_ sites (with different nucleotides between sister groups) would require two substitutions from the parsimony perspective. Thus, from a parsimony perspective, 1/3 of the G_4_ sites evolve at a rate half as fast as the other 2/3 of the sites. Likewise with the likelihood perspective when multiple substitutions are corrected, 2/3 of the G_4_ sites will have a substitution rate at least twice as large as 1/3 of the G_4_ sites, giving rise to rate heterogeneity not present with a star tree. Thus, the relative number of G_4_ and G_3_ sites (denoted n_4_ and n_3_, respectively, Fig. 1A) affect not only the tendency to reject the star tree, but also the characterization of the rate heterogeneity over sites. In what follows, I assess the effect of (n_4_ – n_3_) on the tendency to reject the star tree and the confounding effect of topology on estimating the shape parameter α.

## 2. MATERIALS AND METHODS

### Sets of sequences to make different points

I provide five additional sets of sequences in fasta format as representative of various site pattern combinations. Sample1.fas and Sample2.fas have sequences equivalent to site pattern combinations (0, 14, 43, 4, 40) and (1, 11, 41, 5, 42), respectively (Fig. 1A). They are from sequence simulation under JC69 model with a star tree using Evolver in PAML (Yang, 2007b) to show that likelihood methods will indeed recover a star tree if sequences indeed evolve according to a star tree and a specific substitution model.

Sample3.fas includes all 256 possible site patterns, i.e., all 24 G_1_ sites, 144 G_2_ sites, 48 G_3_ sites, 36 G_4_ sites and 4 G_5_ sites. It is represented by the combination of (24,24,12,12,4). Such site patterns are expected from sequence evolution under JC69 for an infinitely long time so that all sites have experienced full substitution saturation. It is used to highlight the point that the likelihood method correctly recovers the star tree as it should even with full substitution saturation.

Supplemental sequence files Sample4.fas, and Sample5.fas are used to make the main points in the paper. Sample4.fas features a site pattern combination of (24,24,12,96,32) from which likelihood method will not recover a star tree, although the sequences are equidistant from each other with distances computed with any substitution models. As mentioned earlier, this data set is a concatenation of site patterns from simulations with four different trees, i.e., the star tree and the three resolved trees based on JC69 with a constant rate over sites, with slight modifications to ensure that 1) nucleotide frequencies are exactly equal to 0.25, 2) all pairwise comparisons lead to exactly the same number of transitional and transversional differences, and 3) the ratio of transitional and transversional differences between each pairwise comparison is exactly 1/2. Because of the concatenation of sequences simulated from four different topologies, the sequence set represents a special deviation from any of the four topologies (the star tree plus the three resolved trees). While the four sequences will remain equidistant from each other with the site pattern combination of (24,24,12,96,32), the likelihood method will strongly reject the star tree in spite of equidistance among sequences. The sequence set is also used to investigate the confounding effect of tree topology on the estimation of shape parameter α of gamma distribution.

Supplemental file Sample5.fas is used to assess the effect of increasing (n_4_-n_3_) on decreasing support for the star tree. It contains 16 sets of sequences (with each set having fours sequences) of equal sequence length of 2528 derived as follows. A simulation of 100 sets of sequences with a star tree, JC69 model without rate heterogeneity, a sequence length of 2528, and a tree length of 4.8, leads to site pattern combinations that average to (168,216,156,120,80). This site pattern combination, analyzed by either New2.R or NewGamma3.R, leads to the same tree lnL. The continuous gamma model with a star tree yields lnL equal to −13965.18, shape parameter α equal to 5000 (maximum set in optimization), and branch lengths equal to 1.118709. Exactly the same results were obtained with a resolved tree with an additional internal branch length equal to 0. I changed n_3_ and n_4_ values to explore the effect of (n_4_ – n_3_) on the changing support for the star tree and on parameter estimation. In order to maintain the sequence length of 2528, n_1_, n_2_ and n_5_ were also adjusted.

### Likelihood method implemented in R for star tree and gamma-distributed rate

For anyone to recreate results in this paper, I have implemented the likelihood method in R for the JC69 model and four OTUs to evaluate a star tree and a rooted tree, either with a constant rate over sites (Supplemental New2.R) or with a continuous gamma-distributed rate (NewGamma3.R). These are plain text files, self-contained, and well-annotated at the beginning of the file. One can copy and paste them into an R window to obtain results reported here. Computing likelihood with the pruning algorithm has been numerically illustrated in detail (Xia, 2018b).

Because the continuous gamma requires integration over the rate, I used the R function ‘integral’ in the ‘pracma’ package which therefore needs to be installed before running the R scripts. The tree evaluation with a continuous gamma may take three minutes to complete on a PC with an i7-4770 CPU. I also used PhyML (Guindon et al., 2010) to obtain lnL for the resolved tree. I set the tree improvement option ‘-s’ to ‘BEST’ (best of NNI and SPR search), and the ‘-o’ option to ‘tlr’ which optimizes the topology, the branch lengths and rate parameters. For JC69+Γ model, four categories of rates were used to estimate the shape parameter. PhyML does not generate lnL for a star tree, hence the need for the R scripts.

## 3. RESULTS AND DISCUSSION

### Likelihood method recovers the true star tree when sequences evolve under a substitution model

This part, while trivial, is needed for methodological validation. I have simulated sequence evolution of four OTUs under JC69 model and a star tree, by using the Evolver program in the PAML package (Yang, 2007b), with different tree lengths and sequence lengths from 500 to 3000, each with 100 sets of sequences, with a constant rate over site. The sequences are then processed in DAMBE (Xia, 2018a) so that the six site patterns in G_2_ (Fig. 1A) occur exactly equally, so do the four site patterns in G_3_ and three site patterns in G_4_ (Fig. 1A). This ensures that the resulting sequences do not support any one of the three alternative topologies. I analyzed these site patterns by both PhyML 3.1 (Guindon et al., 2010) and the likelihood methods implemented in R in the Supplemental New2.R file and NewGamma3.R.. All these methods recover the star tree, which has the same tree lnL as any of the resolved tree. Furthermore, the branch lengths are equal and nearly identical to the tree length in the input tree used for simulation.

Table 1 includes results for two such simulated sets of sequences without rate heterogeneity (A and B in the column headed by ‘Data’, Table 1). These two sets of sequences are in supplemental Sample1.FAS and Sample2.Fas. Their sequence patterns are represented by the combination of (0,14,43,4,40) and (1,11,41,5,42), respectively, for input to the R scripts. The resolve tree and the star tree consistently have the same tree lnL, whether evaluated by New2.R or PhyML (Table 1). Take Sample1.fas (Data A in Table 1) for example. The resolved tree and the star tree both have lnL of −1537.68, and the branch length for the internal branch of the resolved tree is b5 = 0 (Table 1). PhyML outputs b5 = 0.000001 which is likely the lower bound set for lnL maximization. Analyzing these two sets of sequences with JC69+ Γ by PhyML or my NewGamma3.R script generates the same branch lengths and tree lnL, except for a very large shape parameter α in the order of 10000 (which is expected because these two data sets were simulated with no rate heterogeneity over sites).

**Table 1.** Results of likelihood-based phylogenetic estimation of branch lengths (b_1_ to b_5_ corresponding to x_1_ to x_5_ in Fig. 1B and Fig. 1C) without using gamma-distributed rates to accommodate rate heterogeneity over sites, from running the attached R script (New2.R) and PhyML 3.1 (Guindon et al., 2010).

| Data<sup>(1)</sup> | Method | $b_1$ | $b_2$ | $b_3$ | $b_4$ | $b_5$<sup>(2)</sup> | $\ln L$ |
| --- | --- | --- | --- | --- | --- | --- | --- |
| A | New2.R | 0.426784 | 0.426784 | 0.426784 | 0.426784 | 0.000000 | -1537.68 |
|  |  | 0.426784 | 0.426784 | 0.426784 | 0.426784 | N/A | -1537.68 |
|  | PhyML | 0.426785 | 0.426783 | 0.426793 | 0.426769 | 0.000001 | -1537.68 |
| B | New2.R | 0.400581 | 0.400575 | 0.400584 | 0.400578 | 0.000000 | -1416.09 |
|  |  | 0.400584 | 0.400584 | 0.400584 | 0.400584 | N/A | -1416.09 |
|  | PhyML | 0.400586 | 0.400578 | 0.400570 | 0.400584 | 0.000001 | -1416.09 |
| C | New2.R | 5.655316 | 2.585728 | 10 | 10 | 5.63553 | -1419.57 |
|  |  | 10 | 10 | 9.999938 | 10 | N/A | -1419.57 |
|  | PhyML | 10.000000 | 6.303193 | 10.000000 | 6.180340 | 6.303193 | -1419.57 |

(1) A to C corresponding to supplemental file Sample1.fas to Sample3.fas, respectively, with site pattern combination (0,14,43,4,40), (1,11,41,5,42), and (24,24,12,12,4), respectively.
(2) N/A indicates $b_5$ set to 0 for the star tree.

Data set C (Table 1) is in Sample3.fas with the site pattern combination of (24,24,12,12,4). It represents sequences having experienced full substitution saturation under JC69 model and should have infinitely long branch lengths. However, for practical computation one always set an upper limit for branch lengths, e.g., 10, partly because 1) it is rare for branch lengths to be longer than 10 in practice, and 2) lnL changes little when a branch length increases beyond 10. Minimum and maximum branch lengths can be specified in my two R scripts. Because PhyML 3.1 does not seem to allow branch lengths to go above 10, for comparison I have presented branch lengths and tree lnL with maximum branch lengths set to 10 (Table 1). Both the resolved tree and the star tree have the same lnL of −1419.57. One can change the maximum branch lengths to 100, 1000 or greater in my R scripts, and the star tree, which has one fewer branch to estimate, always has the same lnL as the resolved tree. Thus, the likelihood method does it job well as long as sequences evolve strictly according to substitution model, even with sequences having experienced full substitution saturation.

### Likelihood method fails to recover a star tree with equidistant sequences

The likelihood method rejects a star tree for sequences in supplemental file Sample4.fas, with a site pattern combination of (24,24,12,96,32), in spite of equidistance among the sequences, either with or without gamma-distributed rate (Table 2). When a constant rate is assumed, the tree lnL is −2943.148 for a resolved tree in contrast to −2947.793 for a star tree (Table 2). A likelihood ratio test would lead to 2ΔlnL = 9.29, DF = 1, p = 0.0023 and a rejection of the star tree in favour of a resolved tree. The phylogenetic result with JC69+Γ rejects the star tree more strongly, with 2ΔlnL = 58.03, DF = 1, p = 2.58*10^-14^. Note that the sequences length for this data set is only 536 nt. Longer alignment length would lead to even stronger rejection of the star tree.

**Table 2.** Evaluate two alternative topologies and their branch lengths (b_i_) in Fig. 1B (with b5 set to 0 and not evaluated) and Fig. 1C. Data in supplemental Sample4.fas file, with site pattern combination (24,24,12,96,32).

| Method<sup>(1)</sup> | $b_1$ | $b_2$ | $b_3$ | $b_4$ | $b_5^{(2)}$ | $\alpha^{(3)}$ | lnL |
| --- | --- | --- | --- | --- | --- | --- | --- |
| New2.R | 0.849814 | 0.849814 | 1.693866 | 0 | 0.850 | N/A | -2943.148 |
|  | 1.020276 | 1.020276 | 1.020276 | 1.020276 | N/A | N/A | -2947.793 |
| PhyML | 0.849942 | 0.849764 | 1.694148 | 0.000015 | 0.850 | N/A | -2943.148 |
| NewΓ3 | 2.783640 | 1.133961 | 1.575337 | 2.343977 | 10 | 1 | -2930.116 |
|  | 3.359848 | 1.413517 | 1.886961 | 2.886549 | 30 | 0.843 | -2921.743 |
|  | 3.767304 | 1.850750 | 2.302868 | 3.315211 | 50 | 0.745 | -2918.782 |
|  | 1.020429 | 1.020429 | 1.020429 | 1.020429 | N/A | 10000 | -2947.797 |
| PhyML | 2.213152 | 1.976554 | 2.206997 | 1.968304 | 10 | 0.814 | -2927.806 |

(1) Results from running Supplemental R scripts New2.R and NewGamma3.R (New $\Gamma$ 3), and PhyML.
(2) $b_5$ is $x_5$ in Fig. 1. N/A indicates $b_5$ set to 0 for a star tree.
(3) Shape parameter in gamma distribution, inapplicable (N/A) in phylogenetic reconstruction with a constant rate.

While a site pattern of (24,24,12,96,32) might be considered too extreme, one may take a milder site pattern combination such as (120,192,120,204,164), with a sequence length of 2528. The resulting lnL based on JC69+Γ is −13880.12 for a resolved tree, but −13889.4900 for a star tree, so 2ΔlnL = 18.74. The star tree is rejected with p = 0.000015.

One might argue that there is nothing wrong with the likelihood method itself because the data set is pathological. As mentioned in the Materials and Methods section, this data set is from a concatenation of simulated sequences from four topologies (the star tree and the three resolved trees) with modifications so that nucleotide frequencies are equal and all pairwise comparisons lead to the same number of transition and transversional differences. Thus, although the sequences are equidistant from each other, the star tree is not an appropriate model for the concatenated sequences (neither is any of the three resolved topologies). However, the issue at hand is not on the validity of the likelihood approach, but on its robustness in phylogenetic reconstruction with conflicting phylogenetic signals. With sequences equidistant from each other, we do desire a star tree instead of having it conclusively rejected. Furthermore, we have the problem of inconsistent parameter estimation that I highlight below.

### Inconsistent parameter estimation of likelihood method

The estimated parameter values in Table 2 are disconcerting in two ways. First, when the star tree is imposed, there is no rate heterogeneity over sites, with the shape parameter α in the order of 10000 (Table 2). However, the estimated α becomes 0.745 when b_5_ (x_5_ in Fig. 1) is allowed to be greater than 0, indicating strong rate heterogeneity over sites. I have given reasons in Fig. 2, illustrated with G_4_ sites, that an excess of G_4_ sites will result in rate heterogeneity when a resolved tree is imposed but no rate heterogeneity when a star tree is imposed. That is, given a set of G_4_ sites supporting the three resolved topologies equally, 1/3 of the G_4_ sites will share a low rate of substitution whereas the other 2/3 will share a high rate of substitution when a resolved topology is imposed. In contrast, all G_4_ sites will have the same rate of substitution when a star tree is imposed. This highlights the point that we do need a star tree instead of having three equally supported resolved trees because we need the star tree to perform proper parameter estimation in this case. As I mentioned before, the sequence data is a concatenation of sequence simulation on four different topologies (star and three resolved trees), all with JC69 with a constant rate. However, the maximum likelihood criterion does not allow us to choose the star tree.

Second, the substantial change of lnL with b_5_ under JC69+Γ, associated with a change in α, is also unpleasant. For all practical consideration, the range of values for b_5_ from 10 to 50 all just indicates a branch experiencing substitution saturation, and one would expect lnL to change little with b_5_ in this range of values. However, lnL is −2930.116 with b_5_ = 10 (and α = 1, Table 2), but −0.2918.782 with b_5_ = 50 (and α = 0.745, Table 2). Given that the sequences are equidistant from each other, we do desire a tree with b_5_ = 0, not a method that says that a tree with an unrealistically long b_5_ is the best tree. Note that PhyML (version 3.1) appears to have set the maximum branch length to 10, so its generated lnL (= −2927.806, Table 2) has not yet reached its maximum.

While α is known to depend on topology, phylogenetic researchers often interpret α as if it is a sequence-specific property. A small α is often interpreted to mean strong purifying selection constraining certain sites but not others. I hope that the illustration above will caution against such interpretation.

### The tendency to reject the star tree with increasing (n_4_ − n_3_)

I used sequences in Sample5.fas to assess the effect of (n_4_ − n_3_). The file contains 16 sets of sequences of the same length (=2528) but with different site patterns (n_1_,n_2_,n_3_,n_4_,n_5_), in particular with different (n_4_ − n_3_) values. We may take likelihood chi-square statistic 2ΔlnL as a measure of the tendency to reject the star tree, and expect it to increase with (n_4_ − n_3_) for reasons outlined in Fig. 2. This expectation is consistent with the empirical evidence (Fig. 3). The relationship is stronger between 2ΔlnL and n_4_/n_3_. Also, if we change on n_4_, but keep n_1_, n_2_, n_3_, and n_5_ constant, then 2ΔlnL increases smoothly with (n_4_ − n_3_) or with n_4_/n_3_.

**Fig. 3.**
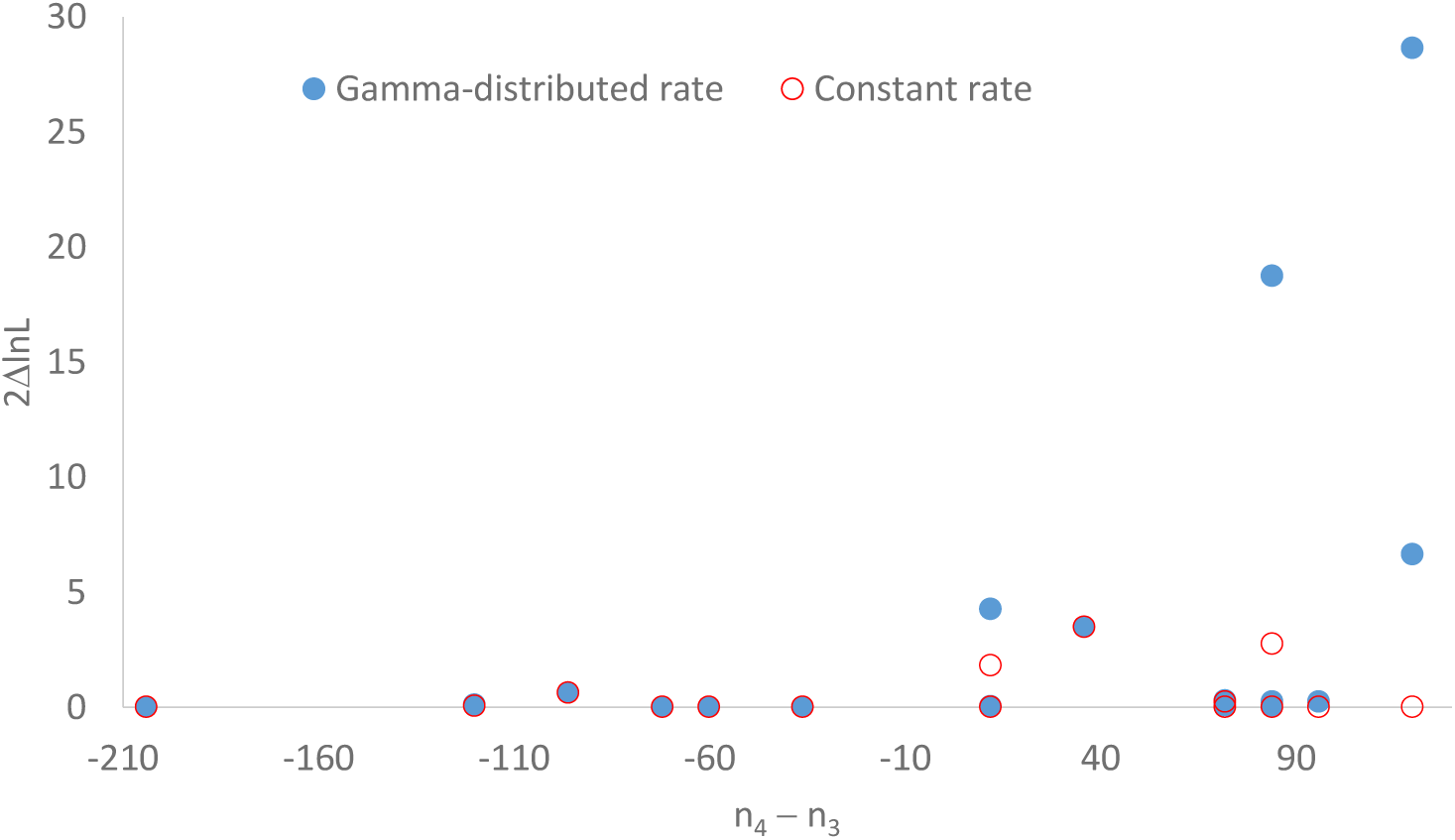
Support for a resolved tree against the star tree, measured by 2ΔlnL = 2(lnL_resolved tree_ − lnL_star tree_), increases with increased G_4_ sites relative to G_3_ sites (measured by n_4_ – n_3_). The open circles indicates the mean (n_4_ – n_3_) value from 100 simulated sequences with a star tree and x_1_ = x_2_ = x_3_ = x_4_ = 1.2.

Fig. 3 indicates that the relationship between 2ΔlnL and (n_4_ – n_3_) is weaker when rate heterogeneity over sites is modelled by a gamma distribution. However, it is not always so, and one can easily find a counter example in which a constant-rate model leads to rejection of the star tree and a gamma-distributed rate model does not. For example, for a site pattern combination of (400,0,0,0,400) we will have lnL = −3984.64 for a resolved tree with JC69, lnL = −3987.998 for a star tree. This leads to 2ΔlnL = 6.716 and a rejection of the star tree (DF = 1, p = 0.0096). With JC69+Γ, we will have lnL = −3546.648 for a resolved tree, lnL = −3547.94 for a star tree. This leads to 2ΔlnL = 2.584 and the star tree is not rejected (DF = 1, p = 0.1079).

This starless bias associated with the ML method sheds light on the well-known star-tree paradox in which Bayesian phylogenetic inference prefers one of the three topologies when the true tree is a star tree (Lewis et al., 2005; Yang, 2007a; Yang and Zhu, 2018). The finding reported here suggests that the problem may not be caused by Bayesian inference but instead is caused by the ML method that favors a resolved tree against the star tree, in particular an excess of G_4_ sites providing conflicting signals supporting three resolved trees equally. It also suggests that the two proposed solutions (Lewis et al., 2005; Yang, 2007a) may not resolve the problem. The reversible-jump Markov chain Monte Carlo algorithm proposed by Lewis et al. (2005), albeit ingenious, is unlikely to solve the problem because the starless bias is not due to the star tree being excluded from tree searching but because it has a smaller likelihood value than any of the three resolved tree. The solution of assigning nonzero prior probability for the degenerate star tree (Lewis et al., 2005; Yang, 2007a) also may not work because the problem is in the likelihood component, not the prior component of Bayesian inference.

I should finally mention that the starless bias and parameter-estimation bias illustrated by the site pattern combination of (24,24,12,96,32) in Sample4.fas is also caused by model misspecification, in the sense that no tree model is appropriate for the data (because it is concatenation of sequences simulated from the star tree and three resolved trees). Existing model-testing methods do not address this type of model misspecification. The conventional selection of the best substitution model will choose JC69 because lnL is the same from JC69 to GTR. This would give us false confidence that we are using an appropriate substitution model for data analysis, without realizing that no tree model is appropriate (i.e., neither the star tree nor the resolved trees) for this set of concatenated sequences. I suggest that, for each phylogeny (T) obtained from a set of sequence data (S) under a substitution model (M), we should obtain the empirical site patterns from S and compare these empirical site patterns against their expectation from sequence simulation based on T and M. If the empirical site patterns deviate significantly from the expectation, then we conclude that the tree model is not appropriate. Applying this approach to sequences in Sample4.fas, we will find the empirical site patterns from the sequence data differ highly significantly from the expectation, and we can conclude that no tree model is appropriate for this set of concatenated sequences or, more specifically, that the sequences are from incompatible tree models.

## ACKNOWLEDGEMENTS

This study is supported by Discovery Grant from Natural Science and Engineering Research Council (NSERC, RGPIN/ 2018-03878) of Canada. I thank Guy Baele, Sudhir Kumar, Laura Kubatko and Arindam RoyChoudhury for discussion. Z. Yang and two anonymous reviewers provided comments that improved clarity of the manuscript.

