## Supplemental for "Starless bias and parameter-estimation bias in the likelihood-based phylogenetic method": New2.R

```
# For 4 sequences, there are 256 possible site patterns. Different combinations of 
#   these site patterns support different substitution models and and differnet
#   topologies.
# For JC69, these 256 site patterns can be reduced to 15 which were boxed into
#   5 site pattern groups (G1 to G5) below. 
#
# 15 site patterns relevant to JC69, into five site pattern groups (G1 to G5):
#    G1   G2     G3   G4  G5
#    1  234567  8901  234  5
# S1 A  TCTGCG  CCGT  ACA  A CGT
# S2 C  TAGATC  AGGT  ATG  A CGT
# S3 T  ACGCAT  ACTT  GCG  A CGT
# S4 G  CTACTG  ACGA  GTA  A CGT
#
# If a set of sequences is made of the 15 sites above, then it is simply represented
#   as a site pattern combination of (1,1,1,1,1). If it is made of 3 G1 site, 24 G2 sites, 
#   8 G3 sites, 12 G4 sites and 12 G5 sites, then it is represented as a site pattern
#   combination of (3,4,2,4,12), i.e., (3/1, 24/6, 8/4, 12/3, 12/1).
#
# Of the 256 possible site patterns for 4 OTUs, 24 site patterns are in G1, 144 in G2, 
#   48 in G3, 36 in G4 and 4 in G5. If a set of sequences happen to be made
#   of these 256 site patterns, then nucleotide frequencies = 0.25, and we observe 
#   exactly 2 transversions to 1 transition (equilibrium expectation). This is also  
#   true for sites within each Group, i.e., the 24 site patterns within G1, or 
#   144 sites patterns in G2, or 48 site patterns in G3, will also have 
#   nuc frequencies of 0.25 and 2 observed transversions to 1 transition
#
# Site patterns within each group jointly support the three alternative topologies
#   equally, i.e., a set of sequences with n1 G1 sites, n2 G2 sites, n3 G3 sites, n4 G4 
#   sites and n5 G5 sites, for a total of (n1*1 + n2*6 + n3*4 + n4*3 + n5*1) sites,
#   will ensure that the 4 sequences are equidistant from each other. In short, the 4
#   sequences are equidistant from each other for any combination of
#   (n1, n2, n3, n4, n5).
# 
# This R script will evaluate two topologies, with the 2nd being the star tree:
# Tree1 ((S1:b1, S2:b2):b5, S3:b3, S4:b4);
# Tree2 ((S1:b1, S2:b2):0, S3:b3, S4:b4); 
#
# Paste the script into an R window, and then enter the equivalent of
#
# sol1<-optim(c(1,1,1,0,1),GetTreeLike,n=c(24,24,12,96,32), method="L-BFGS-B",lower=rep(0,5));
# sol2<-optim(c(1,1,1,1),GetTreeLike,n=c(24,24,12,96,32), method="L-BFGS-B",lower=rep(0,4));
# sol; sol2;
#
# where c(1,1,1,0,1) is the initial value for 5 branch lengths b1 to b5, and 'lower' 
# specifies the lower limits for the five branches, i.e., 0. It seems that 'optim' may not
# explore bi = 0 if none of the initial values is 0. The 'sol1' statement is for the resulved
# Tree1 above, and the 'sol2' statement is for the star tree (with only four branch lengths).
#
# The last few lines are examples (and generate the results in the manuscript).
#
# Given that the 4 sequences are equidistant from each other, we explore the scenarios where 
#   lnL for Tree2 is significantly smaller than lnL for Tree1. The objects sol1 and sol2 each  
#   contains the evaluated branch lengths and lnL for Tree1 and Tree2, respectively (Pay 
#   attention to convergence).
# 
# You can omit one or multiple Groups by specifying 0, e.g., (0,0,0,12,4) will omit site 
#   patterns in Groups 1-3.
#
# For the equivalent implementation with a continuous gamma-distributed rate, see 
#   file NewGamma3.R.
Pii <-  function(x) 1/4+3/4 * exp(-4*x/3);
Pij <-  function(x) 1/4-1/4 * exp(-4*x/3);
arrayA <- c(1,0,0,0);
arrayC <- c(0,1,0,0);
arrayG <- c(0,0,1,0);
arrayT <- c(0,0,0,1);
# L1 and L2 are likelihood vectors for child1 and child2, b1 and b2 are branch lengths to
#   child1 and child2, respectively. The function returns the likelihood vector for the 
#   parent of the two children.
Like <- function(L1,L2,b1,b2) {
  outL <- c(0,0,0,0);
  for(i in 1:4) {
    S1 <- 0; S2 <-0; 
    for(j in 1:4) {
      if(i==j) {
        S1 <- S1+Pii(b1)*L1[j];
        S2 <- S2+Pii(b2)*L2[j];
      } else {
        S1 <- S1+Pij(b1)*L1[j];
        S2 <- S2+Pij(b2)*L2[j];
      }
    }
    outL[i] <- S1*S2;
  }
  return(outL);
}
#---------
# S1           S3
#  \b[1]   b[3]/
#   \_________/
#   / t5   t6 \
#  /b[2]   b[4]\
# S2           S4
# 
# This function returns the likelihood of the 15 site patterns relevant to JC69
# Parameter b is a vector of five branch lengths in the order of b1 to b5.
# For a star tree, the b vector contains b1 to b4, with b5 = 0.
GetSiteL <- function(b) {
  L <- c(rep(0,15));
  if (length(b) == 4) b[5]=0
  # G1 site: A, C, G, T
  arrayX <- Like(arrayA,arrayC,b[1],b[2]);
  arrayY <- Like(arrayG,arrayT,b[3],b[4]);
  arrayZ <- Like(arrayX,arrayY,0,b[5]);
  for(i in 1:4) {
    L[1] <- L[1]+arrayZ[i];
  }
  L[1] <- 0.25*L[1];
  # --------------------------------
  # G2 sites: 2X, Y, Z; L[2] to L[7]
  arrayX <- Like(arrayT,arrayT,b[1],b[2]);
  arrayY <- Like(arrayA,arrayC,b[3],b[4]);
  arrayZ <- Like(arrayX,arrayY,0,b[5]);
  for(i in 1:4) {
    L[2] <- L[2]+arrayZ[i];
  }
  L[2] <- 0.25*L[2];
  #
  arrayX <- Like(arrayC,arrayA,b[1],b[2]);
  arrayY <- Like(arrayC,arrayT,b[3],b[4]);
  arrayZ <- Like(arrayX,arrayY,0,b[5]);
  for(i in 1:4) {
    L[3] <- L[3]+arrayZ[i];
  }
  L[3] <- 0.25*L[3];
  #
  arrayX <- Like(arrayT,arrayG,b[1],b[2]);
  arrayY <- Like(arrayG,arrayA,b[3],b[4]);
  arrayZ <- Like(arrayX,arrayY,0,b[5]);
  for(i in 1:4) {
    L[4] <- L[4]+arrayZ[i];
  }
  L[4] <- 0.25*L[4];
  # 
  arrayX <- Like(arrayG,arrayA,b[1],b[2]);
  arrayY <- Like(arrayC,arrayC,b[3],b[4]);
  arrayZ <- Like(arrayX,arrayY,0,b[5]);
  for(i in 1:4) {
    L[5] <- L[5]+arrayZ[i];
  }
  L[5] <- 0.25*L[5];
  #
  arrayX <- Like(arrayC,arrayT,b[1],b[2]);
  arrayY <- Like(arrayA,arrayT,b[3],b[4]);
  arrayZ <- Like(arrayX,arrayY,0,b[5]);
  for(i in 1:4) {
    L[6] <- L[6]+arrayZ[i];
  }
  L[6] <- 0.25*L[6];
  #
  arrayX <- Like(arrayG,arrayC,b[1],b[2]);
  arrayY <- Like(arrayT,arrayG,b[3],b[4]);
  arrayZ <- Like(arrayX,arrayY,0,b[5]);
  for(i in 1:4) {
    L[7] <- L[7]+arrayZ[i];
  }
  L[7] <- 0.25*L[7];
  # ---------------------------------
  # G3 sites: 3X, Y; L[8] to L[11]
  arrayX <- Like(arrayC,arrayA,b[1],b[2]);
  arrayY <- Like(arrayA,arrayA,b[3],b[4]);
  arrayZ <- Like(arrayX,arrayY,0,b[5]);
  for(i in 1:4) {
    L[8] <- L[8]+arrayZ[i];
  }
  L[8] <- 0.25*L[8];
  #
  arrayX <- Like(arrayC,arrayG,b[1],b[2]);
  arrayY <- Like(arrayC,arrayC,b[3],b[4]);
  arrayZ <- Like(arrayX,arrayY,0,b[5]);
  for(i in 1:4) {
    L[9] <- L[9]+arrayZ[i];
  }
  L[9] <- 0.25*L[9];
  #
  arrayX <- Like(arrayG,arrayG,b[1],b[2]);
  arrayY <- Like(arrayT,arrayG,b[3],b[4]);
  arrayZ <- Like(arrayX,arrayY,0,b[5]);
  for(i in 1:4) {
    L[10] <- L[10]+arrayZ[i];
  }
  L[10] <- 0.25*L[10];
  #
  arrayX <- Like(arrayT,arrayT,b[1],b[2]);
  arrayY <- Like(arrayT,arrayA,b[3],b[4]);
  arrayZ <- Like(arrayX,arrayY,0,b[5]);
  for(i in 1:4) {
    L[11] <- L[11]+arrayZ[i];
  }
  L[11] <- 0.25*L[11];
# ---------------------------------
  # G4 sites: 2X, 2Y; L[12] to L[14]
  arrayX <- Like(arrayA,arrayA,b[1],b[2]);
  arrayY <- Like(arrayG,arrayG,b[3],b[4]);
  arrayZ <- Like(arrayX,arrayY,0,b[5]);
  for(i in 1:4) {
    L[12] <- L[12]+arrayZ[i];
  }
  L[12] <- 0.25*L[12];
  #
  arrayX <- Like(arrayC,arrayT,b[1],b[2]);
  arrayY <- Like(arrayC,arrayT,b[3],b[4]);
  arrayZ <- Like(arrayX,arrayY,0,b[5]);
  for(i in 1:4) {
    L[13] <- L[13]+arrayZ[i];
  }
  L[13] <- 0.25*L[13];
  #
  arrayX <- Like(arrayA,arrayG,b[1],b[2]);
  arrayY <- Like(arrayG,arrayA,b[3],b[4]);
  arrayZ <- Like(arrayX,arrayY,0,b[5]);
  for(i in 1:4) {
    L[14] <- L[14]+arrayZ[i];
  }
  L[14] <- 0.25*L[14];
 
# ----------------------------------
  # G5 site:
  arrayX <- Like(arrayA,arrayA,b[1],b[2]);
  arrayY <- Like(arrayA,arrayA,b[3],b[4]);
  arrayZ <- Like(arrayX,arrayY,0,b[5]);
  for(i in 1:4) {
    L[15] <- L[15]+arrayZ[i];
  }
  L[15] <- 0.25*L[15];
  return(L);   
}
#
# Return the likelihood for the tree given branch lengths and sequences (represented by a 
#   site pattern combination, e.g., (0,14,43,4,40).
GetTreeLike <- function(b,n) {
  L <- GetSiteL(b);
  lnL <- n[1]*log(L[1]);
  for(i in 2:7) {
    lnL <- lnL+n[2]*log(L[i]);
  }
  for(i in 8:11) {
    lnL <- lnL+n[3]*log(L[i]);
  }
  if(n[4]>0) {
    for(i in 12:14) {
      lnL <- lnL+n[4]*log(L[i]);
    }
  }
  if(n[5]>0) {
    lnL <- lnL+n[5]*log(L[15]);
  }
  return(-lnL);
}
# I set initial b4 = 0 below, because otherwise optim may not explore bi = 0 even if lower 
#   bounds are set to 0.
# sol<-optim(c(1,2,1,0,1),GetTreeLike,n=c(24,24,12,12,4), method="L-BFGS-B",lower=rep(0,5));
# sol2<-optim(c(1,2,1,0),GetTreeLike,n=c(24,24,12,12,4), method="L-BFGS-B",lower=rep(0,4));
# R's optim function seems to work faster with different initial branch lengths such as
#  1,2,1,0,1 than with identical initial branch lengths such as 1,1,1,1,1 or 2,2,2,2,2.
#  However, there would seem to be nothing wrong for using 1,1,1,1 for the star tree 
#  because we do expect the same branch lengths.
# Simul1.fas:
sol<-optim(c(1,2,1,0,1),GetTreeLike,n=c(0,14,43,4,40), method="L-BFGS-B",lower=rep(0.0000001,5),upper=rep(2,5));
sol2<-optim(c(1,1,1,1),GetTreeLike,n=c(0,14,43,4,40), method="L-BFGS-B",lower=rep(0.0000001,4),upper=rep(2,4));
# Simul2.fas:
sol3<-optim(c(1,2,1,0,1),GetTreeLike,n=c(1,11,41,5,42), method="L-BFGS-B",lower=rep(0.0000001,5),upper=rep(2,5));
sol4<-optim(c(1,1,1,1),GetTreeLike,n=c(1,11,41,5,42), method="L-BFGS-B",lower=rep(0.0000001,4),upper=rep(2,4));
# The site pattern combination (24,24,12,12,4) means full substitution saturation and infinitely long branches.
#   Limit maximum branch lengths to 10.
sol5<-optim(c(1,2,1,0,1),GetTreeLike,n=c(24,24,12,12,4), method="L-BFGS-B",lower=rep(0.0000001,5),upper=rep(10,5));
sol6<-optim(c(1,1,1,1),GetTreeLike,n=c(24,24,12,12,4), method="L-BFGS-B",lower=rep(0.0000001,4),upper=rep(10,4));
sol7<-optim(c(1,2,1,0,1),GetTreeLike,n=c(24,24,12,96,32), method="L-BFGS-B",lower=rep(0.0000001,5),upper=rep(10,5));
sol8<-optim(c(1,1,1,1),GetTreeLike,n=c(24,24,12,96,32), method="L-BFGS-B",lower=rep(0.0000001,4),upper=rep(10,4));
```
