## Supplemental for "Starless bias and parameter-estimation bias in the likelihood-based phylogenetic method": NewGamma3.R

```
# Need R library pracma for integration in continous gamma
# Before pasting all into an R window, type 'install.packages("pracma")' if it 
#   has not been done before
# R's optim function needs reasonable initial values to converge. In the last case,
#   computing lnL for a range of branch lengths and alpha and plot3D helps.
#
# For 4 sequences, there are 256 possible site patterns. Different combinations of 
#   these site patterns support different substitution models and and differnet
#   topologies.
# For JC69, these 256 site patterns can be reduced to 15 which were boxed into
#   5 site pattern groups (G1 to G5) below. 
#
# 15 site patterns relevant to JC69, into five site pattern groups (G1 to G5):
#    G1   G2     G3   G4  G5
#    1  234567  8901  234  5
# S1 A  TCTGCG  CCGT  ACA  A CGT
# S2 C  TAGATC  AGGT  ATG  A CGT
# S3 T  ACGCAT  ACTT  GCG  A CGT
# S4 G  CTACTG  ACGA  GTA  A CGT
#
# If a set of sequences is made of the 15 sites above, then it is simply represented
#   as a site pattern combination of (1,1,1,1,1). If it is made of 3 G1 site, 24 G2 sites, 
#   8 G3 sites, 12 G4 sites and 12 G5 sites, then it is represented as a site pattern
#   combination of (3,4,2,4,12), i.e., (3/1, 24/6, 8/4, 12/3, 12/1).
#
# Of the 256 possible site patterns for 4 OTUs, 24 site patterns are in G1, 144 in G2, 
#   48 in G3, 36 in G4 and 4 in G5. If a set of sequences happen to be made
#   of these 256 site patterns, then nucleotide frequencies = 0.25, and we observe 
#   exactly 2 transversions to 1 transition (equilibrium expectation). This is also  
#   true for sites within each Group, i.e., the 24 site patterns within G1, or 
#   144 sites patterns in G2, or 48 site patterns in G3, will also have 
#   nuc frequencies of 0.25 and 2 observed transversions to 1 transition
#
# This R script will evaluate two topologies, with the 2nd being the star tree:
# Tree1 ((S1:b1, S2:b2):b5, S3:b3, S4:b4);
# Tree2 ((S1:b1, S2:b2):0, S3:b3, S4:b4); 
#
# The obtain branch length estimates, alpha parameter, lnL, issue the following:
#
# siteCombo <- c(0,14,43,4,40);
# initialVal <- c(1,0.1,1,3,0.1,5); 
# sol1<-optim(initialVal,GetTreeLike,n=siteCombo, method="L-BFGS-B",lower=c(rep(0.0000001,5),1),upper=c(rep(10,5),20));
# initialVal <- c(0.4,0.4,0.4,0.4,10); 
# sol2<-optim(initialVal,GetTreeLike,n=siteCombo, method="L-BFGS-B",lower=c(rep(0.0000001,4),1),upper=c(rep(10,4),20));
#
# where sol1 is for resolved tree and sol2 for the star tree. The last parameter in 'initialVal' and in 'lower' and
#   'upper' is shape parameter alpha, others being branch lengths (5 for resolved tree and 4 for star tree).
#
# Given that the 4 sequences are equidistant from each other, we explore the scenarios where 
#   lnL for Tree2 is significantly smaller than lnL for Tree1. The objects sol1 and sol2 each  
#   contains the evaluated branch lengths and lnL for Tree1 and Tree2, respectively (Pay 
#   attention to convergence).
# 
# You can omit one or multiple Groups by specifying 0, e.g., (0,0,0,12,4) will omit site 
#   patterns in Groups 1-3.
#
Pii <-  function(x) 1/4+3/4 * exp(-4*x/3);
Pij <-  function(x) 1/4-1/4 * exp(-4*x/3);
arrayA <- c(1,0,0,0);
arrayC <- c(0,1,0,0);
arrayG <- c(0,0,1,0);
arrayT <- c(0,0,0,1);
Like <- function(L1,L2,b1,b2) {
  outL <- c(0,0,0,0);
  for(i in 1:4) {
    S1 <- 0; S2 <-0; 
    for(j in 1:4) {
      if(i==j) {
        S1 <- S1+Pii(b1)*L1[j];
        S2 <- S2+Pii(b2)*L2[j];
      } else {
        S1 <- S1+Pij(b1)*L1[j];
        S2 <- S2+Pij(b2)*L2[j];
      }
    }
    outL[i] <- S1*S2;
  }
  return(outL);
}
#
# The calling function will have b<-c(b1,b2,b3,b4,b5); ACGT0123 <- c(0,1,2,3) for a site ACGT
# SiteLike is the integrand for gamma-distributed model
SiteLike <- function(arrayR,b,alpha,ACGT0123) {
  r.n <- length(arrayR);
  siteL <- c(rep(0,r.n));
  for(i in 1:r.n) {
    r <- arrayR[i]; 
    if (ACGT0123[1]==0) {
      if(ACGT0123[2]==0) {
        arrayX<-Like(arrayA,arrayA,r*b[1],r*b[2]) 
      } else if(ACGT0123[2]==1) {
        arrayX<-Like(arrayA,arrayC,r*b[1],r*b[2])
      } else if(ACGT0123[2]==2) {
        arrayX<-Like(arrayA,arrayG,r*b[1],r*b[2])
      } else {
        arrayX<-Like(arrayA,arrayT,r*b[1],r*b[2])
      }
    } else if(ACGT0123[1]==1) {
      if(ACGT0123[2]==0) {
        arrayX<-Like(arrayC,arrayA,r*b[1],r*b[2])
      } else if(ACGT0123[2]==1) {
        arrayX<-Like(arrayC,arrayC,r*b[1],r*b[2])
      } else if(ACGT0123[2]==2) {
        arrayX<-Like(arrayC,arrayG,r*b[1],r*b[2])
      } else {
        arrayX<-Like(arrayC,arrayT,r*b[1],r*b[2])
      }
    } else if(ACGT0123[1]==2) {
      if(ACGT0123[2]==0) {
        arrayX<-Like(arrayG,arrayA,r*b[1],r*b[2])
      } else if(ACGT0123[2]==1) {
        arrayX<-Like(arrayG,arrayC,r*b[1],r*b[2])
      } else if(ACGT0123[2]==2) {
        arrayX<-Like(arrayG,arrayG,r*b[1],r*b[2])
      } else {
        arrayX<-Like(arrayG,arrayT,r*b[1],r*b[2])
      }
    } else {
      if(ACGT0123[2]==0) {
        arrayX<-Like(arrayT,arrayA,r*b[1],r*b[2])
      } else if(ACGT0123[2]==1) {
        arrayX<-Like(arrayT,arrayC,r*b[1],r*b[2])
      } else if(ACGT0123[2]==2) {
        arrayX<-Like(arrayT,arrayG,r*b[1],r*b[2])
      } else {
        arrayX<-Like(arrayT,arrayT,r*b[1],r*b[2])
      }
    }
  #
    if (ACGT0123[3]==0) {
      if(ACGT0123[4]==0) {
        arrayY<-Like(arrayA,arrayA,r*b[3],r*b[4])
      } else if(ACGT0123[4]==1) {
        arrayY<-Like(arrayA,arrayC,r*b[3],r*b[4])
      } else if(ACGT0123[4]==2) {
        arrayY<-Like(arrayA,arrayG,r*b[3],r*b[4])
      } else {
        arrayY<-Like(arrayA,arrayT,r*b[3],r*b[4])
      }
    } else if(ACGT0123[3]==1) {
      if(ACGT0123[4]==0) {
        arrayY<-Like(arrayC,arrayA,r*b[3],r*b[4])
      } else if(ACGT0123[4]==1) {
        arrayY<-Like(arrayC,arrayC,r*b[3],r*b[4])
      } else if(ACGT0123[4]==2) {
        arrayY<-Like(arrayC,arrayG,r*b[3],r*b[4])
      } else {
        arrayY<-Like(arrayC,arrayT,r*b[3],r*b[4])
      }
    } else if(ACGT0123[3]==2) {
      if(ACGT0123[4]==0) {
        arrayY<-Like(arrayG,arrayA,r*b[3],r*b[4])
      } else if(ACGT0123[4]==1) {
        arrayY<-Like(arrayG,arrayC,r*b[3],r*b[4])
      } else if(ACGT0123[4]==2) {
        arrayY<-Like(arrayG,arrayG,r*b[3],r*b[4])
      } else {
        arrayY<-Like(arrayG,arrayT,r*b[3],r*b[4])
      }
    } else {
      if(ACGT0123[4]==0) {
        arrayY<-Like(arrayT,arrayA,r*b[3],r*b[4])
      } else if(ACGT0123[4]==1) {
        arrayY<-Like(arrayT,arrayC,r*b[3],r*b[4])
      } else if(ACGT0123[4]==2) {
        arrayY<-Like(arrayT,arrayG,r*b[3],r*b[4])
      } else {
        arrayY<-Like(arrayT,arrayT,r*b[3],r*b[4])
      }
    }
    arrayZ <- Like(arrayX,arrayY,0,r*b[5]);
    for(j in 1:4) {
      siteL[i] <- siteL[i]+arrayZ[j];
    }
    siteL[i] <- 0.25*siteL[i]*dgamma(r,shape=alpha,scale=1/alpha);
  }
  return(siteL);
}
#---------
# N1           N3
#  \b[1]       b[3]/
#   \_________/
#   / t5   t6 \
#  /b[2]       b[4]\
# N2           N4
#   1 2   3 4
# ((A,C),(T,G))
# A, C, G, T
# 15 site patterns (see top) in 5 groups to ensure equidistance among the 4 OTUs
# L1: site 1; L2: sites 2-7; L3: sites 8-11; L4: sites 12-14; L5: site 15
# Use SiteLike <- function(r,b,alpha,ACGT0123) 
GetSiteL <- function(b,alpha) {
  library(pracma);  
  # Ideally, the integration below should have minR = 0 and maxR = Inf, but integral 
  #   does not seem to work consistently with Inf (The same statement may work fine
  #   at one time but not in the other, hence the compromise of defining a finite maxR.
  #   maxR = 1200 should be sufficient for small alpha as alpha unlikely < 0.1.
  # The occasional error that is not informative: 
  # Error in .gkadpt(f, a, (a + b)/2, tol = tol) : object 'Q2' not found
  minR <- 0;
  if(alpha>100) {
    maxR <- 10;
  } else if(alpha>10) {
    maxR <- 15;
  } else if(alpha>4) {
    maxR <- 40;
  } else if(alpha>1) {
    maxR <- 150;
  } else if(alpha>0.5) {
    maxR <- 250;
  } else {
    maxR <- 1200; 
  }

  L <- c(rep(0,15));
  if (length(b) == 4) b[5]=0
  # A, C, G, T
  L[1] <- integral(SiteLike,xmin=minR,xmax=maxR,b=b,alpha=alpha,ACGT0123=c(0,1,2,3));
  # --------------------------------
  # 2X, Y, Z
  L[2] <- integral(SiteLike,xmin=minR,xmax=maxR,b=b,alpha=alpha,ACGT0123=c(3,3,0,1));
  L[3] <- integral(SiteLike,xmin=minR,xmax=maxR,b=b,alpha=alpha,ACGT0123=c(1,0,1,3));
  L[4] <- integral(SiteLike,xmin=minR,xmax=maxR,b=b,alpha=alpha,ACGT0123=c(3,2,2,0));
  L[5] <- integral(SiteLike,xmin=minR,xmax=maxR,b=b,alpha=alpha,ACGT0123=c(2,0,1,1));
  L[6] <- integral(SiteLike,xmin=minR,xmax=maxR,b=b,alpha=alpha,ACGT0123=c(1,3,0,3));
  L[7] <- integral(SiteLike,xmin=minR,xmax=maxR,b=b,alpha=alpha,ACGT0123=c(2,1,3,2));
  # ---------------------------------
  # 3X, Y
  L[8] <- integral(SiteLike,xmin=minR,xmax=maxR,b=b,alpha=alpha,ACGT0123=c(1,0,0,0));
  L[9] <- integral(SiteLike,xmin=minR,xmax=maxR,b=b,alpha=alpha,ACGT0123=c(1,2,1,1));
  L[10] <- integral(SiteLike,xmin=minR,xmax=maxR,b=b,alpha=alpha,ACGT0123=c(2,2,3,2));
  L[11] <- integral(SiteLike,xmin=minR,xmax=maxR,b=b,alpha=alpha,ACGT0123=c(3,3,3,0));
# ---------------------------------
   # 2X, 2Y
  L[12] <- integral(SiteLike,xmin=minR,xmax=maxR,b=b,alpha=alpha,ACGT0123=c(0,0,2,2));
  L[13] <- integral(SiteLike,xmin=minR,xmax=maxR,b=b,alpha=alpha,ACGT0123=c(1,3,1,3));
  L[14] <- integral(SiteLike,xmin=minR,xmax=maxR,b=b,alpha=alpha,ACGT0123=c(0,2,2,0));
# ----------------------------------
  # AAAA
  L[15] <- integral(SiteLike,xmin=minR,xmax=maxR,b=b,alpha=alpha,ACGT0123=c(0,0,0,0));
  return(L);   
}

# Test function to be deleted
TestGetTreeLike <- function(b,alpha,n) {
  x <- c(b,b,b,b);
  L <- GetSiteL(x,alpha);
  lnL <- n[1]*log(L[1]);
  if(n[2]>0) {
    for(i in 2:7) {
      lnL <- lnL+n[2]*log(L[i]);
    }
  }
  if(n[3]>0) {
    for(i in 8:11) {
      lnL <- lnL+n[3]*log(L[i]);
    }
  }
  if(n[4]>0) {
    for(i in 12:14) {
      lnL <- lnL+n[4]*log(L[i]);
    }
  }
  if(n[5]>0) {
    lnL <- lnL+n[5]*log(L[15]);
  }
  return(-lnL);
}

# The last element in vector b is alpha
GetTreeLike <- function(b,n) {
  b.n<-length(b)
  alpha<-b[b.n]
  x<-b[1:(b.n-1)]
  L <- GetSiteL(x,alpha);
  lnL <- n[1]*log(L[1]);
  if(n[2]>0) {
    for(i in 2:7) {
      lnL <- lnL+n[2]*log(L[i]);
    }
  }
  if(n[3]>0) {
    for(i in 8:11) {
      lnL <- lnL+n[3]*log(L[i]);
    }
  }
  if(n[4]>0) {
    for(i in 12:14) {
      lnL <- lnL+n[4]*log(L[i]);
    }
  }
  if(n[5]>0) {
    lnL <- lnL+n[5]*log(L[15]);
  }
  return(-lnL);
}
# 24,24,12,12,4: all 256 possible site patterns for 4 OTUs 
# Set initial values. The last value is alpha (shape parameter for gamma distribution). 
# The upper limit for branch lengths were set to 50 (The upper limit is 4 in PAML's baselml/basemlg, and 10
#   in PhyML) to reduce computation time, but they do not obtain the maximized lnL in some cases. 
# Note: This causes problems in model-test programs such as jModelTest that uses, say, PhyML for comuting lnL.
#   For example, if true maximized lnL is -1000 for JC69 but reported lnL is -1500 because of the branch lengths 
#   were constrained, and if the reported lnL for K80 is -1400, then K80 will be reported as a better model
#   than JC69, which is wrong. However, setting a very large maximum for branch lengths takes longer
#   computation (the continuous gamma is very slow).
# Uncomment the following to evaluate the resolved and the star trees (takes 1-5 min for each evaluation):
# 
# lnL may increase slightly if running twice, i.e., using the output parameter values as initial values in the 2nd run.
#
siteCombo <- c(0,14,43,4,40);
initialVal <- c(1,0.1,1,3,0.1,5); 
sol1<-optim(initialVal,GetTreeLike,n=siteCombo, method="L-BFGS-B",lower=c(rep(0.0000001,5),1),upper=c(rep(10,5),20));
sol3<-optim(initialVal,GetTreeLike,n=siteCombo, method="L-BFGS-B",lower=c(rep(0.0000001,5),1),upper=c(rep(10,5),200));
initialVal <- c(0.4,0.4,0.4,0.4,10); 
sol2<-optim(initialVal,GetTreeLike,n=siteCombo, method="L-BFGS-B",lower=c(rep(0.0000001,4),1),upper=c(rep(10,4),20));
sol4<-optim(initialVal,GetTreeLike,n=siteCombo, method="L-BFGS-B",lower=c(rep(0.0000001,4),1),upper=c(rep(10,4),200));
#
siteCombo <- c(1,11,41,5,42);
initialVal <- c(1,0.1,1,3,0.1,5); 
sol11<-optim(initialVal,GetTreeLike,n=siteCombo, method="L-BFGS-B",lower=c(rep(0.0000001,5),1),upper=c(rep(10,5),20));
sol13<-optim(initialVal,GetTreeLike,n=siteCombo, method="L-BFGS-B",lower=c(rep(0.0000001,5),1),upper=c(rep(10,5),200));
initialVal <- c(0.4,0.4,0.4,0.4,5); 
sol12<-optim(initialVal,GetTreeLike,n=siteCombo, method="L-BFGS-B",lower=c(rep(0.0000001,4),1),upper=c(rep(10,4),20));
sol14<-optim(initialVal,GetTreeLike,n=siteCombo, method="L-BFGS-B",lower=c(rep(0.0000001,4),1),upper=c(rep(10,4),200));
#
siteCombo <- c(24,24,12,12,4);
initialVal <- c(5,8,5,8,9,5); 
sol21<-optim(initialVal,GetTreeLike,n=siteCombo, method="L-BFGS-B",lower=c(rep(0.0000001,5),0.1),upper=c(rep(10,5),20));
sol23<-optim(initialVal,GetTreeLike,n=siteCombo, method="L-BFGS-B",lower=c(rep(0.0000001,5),0.1),upper=c(rep(10,5),200));
initialVal <- c(5,5,5,5,5); 
sol22<-optim(initialVal,GetTreeLike,n=siteCombo, method="L-BFGS-B",lower=c(rep(0.0000001,4),0.1),upper=c(rep(10,4),20));
sol24<-optim(initialVal,GetTreeLike,n=siteCombo, method="L-BFGS-B",lower=c(rep(0.0000001,4),0.1),upper=c(rep(10,4),200));
#
siteCombo <- c(24,24,12,96,32);
initialVal <- c(3,1,1,2,10,0.5); 
sol31<-optim(initialVal,GetTreeLike,n=siteCombo, method="L-BFGS-B",lower=c(rep(0.0000001,5),0.1),upper=c(rep(10,5),1));
sol33<-optim(initialVal,GetTreeLike,n=siteCombo, method="L-BFGS-B",lower=c(rep(0.0000001,5),0.1),upper=c(rep(30,5),2));
sol35<-optim(initialVal,GetTreeLike,n=siteCombo, method="L-BFGS-B",lower=c(rep(0.0000001,5),0.1),upper=c(rep(50,5),2));
# 
# Just to get a rough idea about the parameter space that would maximize lnL for the star tree:
# ind<-10*21;b<-rep(0,ind);alpha<-rep(0,ind);lnL<-rep(0,ind);count<-1;
# for(i in seq(from=1,by=1,to=10)) {
#   for(j in seq(from=1,by=10,to=201)) {
#     b[count] <- i; alpha[count] <- j;
#     lnL[count] <- -TestGetTreeLike(b=i,alpha=j,n=siteCombo);
#     count <- count+1;
#   }
# }
# # plot3D to find the lnL peak
# library(plot3D);
# scatter3D(b,alpha,lnL,labels,pch=20,cex=2,ticktype="detailed",xlab="b",ylab="alpha",zlab="lnL",theta=40,phi=40)
# # max lnL when b = 1 and alpha = 201
# initialVal <- c(1,1,1,1,10000); 
# sol32<-optim(initialVal,GetTreeLike,n=siteCombo, method="L-BFGS-B",lower=c(rep(0.5,4),201),upper=c(rep(1.5,4),10000));
```
